# All-Diamond Boron-Doped Microelectrodes for Neurochemical Sensing with Fast-Scan Cyclic Voltammetry

**DOI:** 10.1101/2024.08.07.606919

**Authors:** Bhavna Gupta, Brandon Kepros, Jann B. Landgraf, Michael F. Becker, Wen Li, Erin K. Purcell, James R. Siegenthaler

## Abstract

Neurochemical sensing with implantable devices has gained remarkable attention over the last few decades. A promising area of this research is the progress of novel electrodes as electrochemical tools for neurotransmitter detection in the brain. The boron-doped diamond (BDD) electrode is one such candidate that previously has been reported for its excellent electrochemical properties, including a wide working potential, superior chemical inertness and mechanical stability, good biocompatibility and resistance to fouling. Meanwhile, limited research has been conducted on the BDD as a microelectrode for neurochemical detection. Our team has developed a freestanding, all diamond microelectrode consisting of a boron-doped polycrystalline diamond core, encapsulated in an insulating polycrystalline diamond shell, with a cleaved planar tip for electrochemical sensing. This all-diamond electrode is advantageous due to its – (1) batch fabrication using wafer technology that eliminates traditional hand fabrication errors and inconsistencies, (2) absence of metal-based wires, or foundations, to improve biocompatibility and flexibility, and (3) sp^3^ carbon surface with resistance to biofouling, i.e. adsorption of proteins or unwanted molecules at the electrode surface in a biological environment that impedes overall electrode performance. Here, we provide findings on further in vitro testing and development of the freestanding boron-doped diamond microelectrode (BDDME) for neurotransmitter detection using fast scan cyclic voltammetry (FSCV). In this report, we elaborate on – 1) an updated fabrication scheme and work flow to generate all diamond BDDMEs, 2) slow scan cyclic voltammetry measurements of reference and target analytes to understand basic electrochemical behavior of the electrode, and 3) FSCV characterization of common neurotransmitters, and overall favorability of serotonin (5-HT) detection. The BDDME showed a 2-fold increased FSCV response for 5-HT in comparison to dopamine (DA), with a limit of detection of 0.16 µM for 5-HT and 0.26 µM for DA. These results are intended to expand on the development of the next generation BDDME and guide future in vivo experiments, adding to the growing body of literature on implantable devices for neurochemical sensing.

## Introduction

Implantable electrodes that can be integrated with biological tissue to monitor and deliver electrical or chemical signals have gained increasing interest in the study of neurological disorders^1–4^. Such microelectrodes and microelectrode arrays have allowed for neural stimulation as well as acute and chronic recordings to better understand, diagnose and treat diseases such as epilepsy, Parkinson’s disease, and depression^5–7^. Recent advances in electrode development have largely focused on electrophysiological, optical, stimulating or hybrid approaches to study electrical activity in the brain^8–10^. In contrast, current progress with sensors to examine real-time and long-term chemical signaling in living systems is relatively limited and challenging^2^. Electrochemical sensors targeted to detect neurotransmitters in physiological environments, such as the brain, face major issues including selective identification of analytes, long-term electrode stability, sparse distribution of neural circuitries and signal loss due to protein adsorption related surface fouling, i.e. biofouling^11–14^.

Exploration of new electrode materials is one avenue towards addressing these challenges and contributing to the next generation of implantable electrochemical tools^15–17^. The carbon-fiber microelectrode (CFME) is the present-day standard implantable electrode for neurochemical sensing with fast-scan cyclic voltammetry (FSCV), a popular technique offering adequate spatial and temporal resolution to study sub-second neurotransmitter release in biological environments. In general, FSCV facilitates redox reactions of analytes by applying a fast-repeating voltage-driven waveform at the microelectrode to measure the resulting change in current. The CFME offers advantageous sp^2^ carbon structure to favor adsorption-controlled processes at the electrode surface maintains the sensitivity to detect neurotransmitters with FSCV. Despite 30 years of use and development, the CFME still faces major drawbacks that limit it as an ideal implantable neurochemical sensor for clinical application. First, the high adsorptive CFME surface in biological conditions is prone to biofouling and degradation resulting in signal sensitivity loss over time. Second, the CFME is conventionally hand fabricated using low-cost assembly methods which produce electrodes that are susceptible to human error, inconsistent in size and limited in shape and structure. As researchers move towards arrays and away from singular fiber fabrication, there has been great development with individual CFMEs being placed on an array and insulated with parylene-c^18,19^. However, these arrays are still hand fabricated. Other FSCV electrode array developments have focused on pyrolyzed photoresist films on a silicon wafer^20^ creating a 4-channel recording probe, glassy carbon microelectrode arrays on a flexible polymer substrate^21,22^ creating multichannel (8+) recording sites. Higher flexibility arrays have been developed using carbon fiber electrode threads^23^ as well as a NeuroString, with a graphene based biosensing neural interface^24^. Most recently has been the development of a fuzzy graphene microelectrode array^25^.

Our team has previously reported on a promising all-diamond boron-doped diamond microelectrode (BDDME) for electrochemical measurements^12,15,26,27^. The boron-doped diamond (BDD) electrode, offering sp^3^ carbon structure, has gained attention as a favorable biosensor due to its low background current, chemical inertness, mechanical stability, large working potential window and biocompatibility^15,28,29^. Others have shown that hybridizing BDD to CFME’s reduces chemical fouling, and increases CFME lifetime^30,31^. Polycrystalline BDD films on sharpened tungsten rods have shown reduced fouling and increased mechanical strength compared to CFME with FSCV measurements. However, BDD electrodes grown on metals such as tungsten or platinum still require hand fabrication and packaging in an insulating medium, and are not easily amenable towards batch electrode array fabrication.

Our team’s all-diamond BDDME is freestanding and contains no additional metals to potentially address the outlined challenges faced by the standard CFME and typical metal-based BDD electrodes. These BDDMEs were constructed using batch wafer technology which is amenable to reproducibility, as opposed to hand fabrication techniques, where all the sensors are processed identically at a time. The sp^3^ to sp^2^ ratio of the all-diamond BDD is tunable during the diamond growth process. The inherent sp^3^ carbon structure of diamond provides a less adsorptive electrode surface, which is advantageous for tackling biofouling, albeit at the expense of sensitivity. Batch wafer fabrication of electrodes produces consistent and closely replicated electrodes that could potentially be used in a microelectrode array. From a commercialization standpoint, modern wafer technology allows for easy scale-up and batch fabrication of electrodes which surpasses conventional hand fabrication techniques producing one sensor at a time. Our recent study on the in vitro biofouling performance^12^ of these BDDMEs suggested that: 1) possible roles of both diffusion-as well as adsorption-controlled processes facilitating neurotransmitter sensing at the BDDME with FSCV, and 2) reduced biofouling-induced current attenuation at BDDME with a waveform selective for a serotonin redox reaction. A deeper investigation of the electrochemical behavior of BDDME was warranted to further explore the findings in this previous work.

In this report, we provide detailed findings on: (1) the fabrication scheme and changes implemented to improve the electrochemical performance of the BDDME, (2) slow scan cyclic voltammetric responses at the BDDME of reference analytes, such as Ruthenium Hexamine and Ferrocene Carboxylic Acid, and target analytes, such as dopamine (DA) and serotonin (5-HT), (3) fast scan cyclic voltammetric responses to common neurotransmitters (DA, 5-HT, 3,4-Dihydroxyphenylacetic acid (DOPAC), Ascorbic Acid (AA) and, hydrogen peroxide (H_2_O_2_) using a custom flow cell injection system and, (4) In vitro FSCV responses to mixture solutions containing varying concentrations of DA and 5-HT for oxidative peak identification. The limit of detection of DA was 0.26 µM and 0.16 µM for 5-HT, calculated using the linear dynamic ranges of each. Overall, the BDDME demonstrated favorability to detect 5-HT over DA, with the 5-HT oxidative peak current response being around two times greater than that of DA for the same concentrations. This report provides new findings to extend upon our work to develop the next-generation BDDME as a chronic, in vivo neurochemical sensor. The findings of this study add to the growing body of literature for next-generation electrochemical sensors and implantable neural devices.

## Experimental Methods

### Chemicals

All chemicals were purchased from Sigma Aldrich (St. Louis, MO, USA) unless otherwise specified. Dopamine (DA), serotonin (5-HT), ascorbic acid (AA), 3,4-Dihydroxyphenylacetic acid (DOPAC) and hydrogen peroxide were all purchased from Sigma-Aldrich, Inc. A 1 µM stock solution of each was prepared in 1 mM perchloric acid (HClO_4_), and final working solutions were prepared diluting the stock solution in pH 7.4 Tris artificial cerebral-spinal fluid (aCSF) buffer immediately before measurement. Calibration ranges were from 200 nM – 200 µM. Tris aCSF was prepared by making a solution of 25 mM Tris, 126 mM NaCl, 2.5 mM KCl, 1.2 mM NaH_2_PO_4,_ 2.4 mM CaCl_2_, 1.2 mM MgCl_2_ and adjusting the solution to pH 7.4^32^. Solution working time for the stocks was kept to no greater than 30 min to prevent measurable differences through degradation.

### Instrumentation/Data Acquisition

FSCV experiments were conducted using a two-electrode setup vs a Ag/AgCl quasi reference electrode. Electrodes were connected to a mini-UEI potentiostat with a variable gain headstage. (UNC Electronics Facility, University of North Carolina, Chapel Hill, NC). Data was collected using an NI-6363 data acquisition card, using HDCV Analysis software (Department of Chemistry, University of North Carolina, Chapel Hill, NC)^33^. Unless otherwise specified, a standard triangular FSCV waveform was used, by applying a -0.4 V holding potential, and ramping to a switching potential of 1.3 V at 400 V/s, repeating at 10 Hz to the electrode. Electrodes were tested and calibrated using a flow cell injection system with a TTL controlled switching source and a flowrate of 0.5 mL/min supplied by a NE-1000 syringe pump (New Era Pump Systems, Inc. Farmingdale, NY). Reference electrodes used in FSCV experiments were fabricated using quasi-Ag/AgCl wires. Quasi-Ag/AgCl electrode were fabricated by soaking a silver wire (0.25 mm, Alfa Aesar) in undiluted concentrated bleach overnight (Clorox Professional Products Company, Oakland, CA).Slow-scan voltammetric (< 1 V/s) and electrical impedance spectroscopy measurements were made using a CHI 660C (CH Instruments, Austin TX) potentiostat with a Pt counter electrode and a Ag/AgCl reference electrode.

### Statistical Methods

Data were analyzed and plotted using Origin (Pro), 2019. OriginLab Corporation, Northampton, MA, USA and Graphpad Prism version 10.3.0 for Windows, GraphPad Software, Boston Massachusetts, USA. Means and standard errors are tabulated and reported. Similar to our previous report, limit of detections (LOD) were calculated from the linear best fit equation with the corresponding sensitivity for the linear dynamic range of each analyte^34^.

### Boron-Doped Diamond Microelectrode (BDDME) fabrication

The BDDME was designed to have an 8 mm long shank, with both a 30 or a 50 µm wide tip, with a larger connection pad for handling and electrical connection. The conductive diamond was designed to be grown 4 µm thick, and insulated with a 10µm layer with a a grow around method. Using a multi-step fabrication process (**Figure 1A**) modified from our previous work, we further developed and optimized these electrodes^27,35^. Previously, the diamond wafers were fabricated in a 2.45 GHz microwave reactor, grown on 3” silicon substrates^27^. In an effort to move towards scalability, 4” ∅ 500 µm thick single side polished silicon wafers were utilized with larger diamond reactors. Initially, wafers were scratch seeded and microcrystalline BDD films were grown on a 4” silicon wafer using a 915 MHz microwave plasma-assisted chemical vapor deposition reactor. Standard BDD synthesis conditions included a microwave power of 9 kW, 900 °C stage temperature, a chamber pressure of 60 Torr and a gas chemistry of 2 % methane. Diborane was added to the diamond grown at a B/C ratio of 37,500 ppm to ensure conductivity. Following BDD growth, copper and titanium were electron beam evaporated (Kurt Lesker Axiss PVD System) and patterned via photolithography (ABM-USA, INC., Jan Jose, CA, USA), (**Figure 1B**) followed by a wet chemical etch. The patterned BDD films with the copper mask were then etched using a microwave assisted reactive ion etcher (RIE) SF_6_/Ar/O_2_ with a microwave power of 1000 W and a radio-frequency (RF) bias of 150 W (180 V). The copper mask was then removed using nitric acid. The Si was removed using an HNA etchant, at a ratio of 6:11:5 acetic acid: nitric acid: hydrofluoric acid. Prior to Si etching, batches of electrodes were laser cut out of the main wafer to decrease etchant time, and allow for ease of handleability. The now released electrodes were then placed on a silicon holder vertically, where an insulting PCD film was grown around the freestanding BDD cores, using a hot filament chemical vapor deposition (HF-CVD) around the BDD fibers, insulating the conductive diamond.

**Figure 1:**
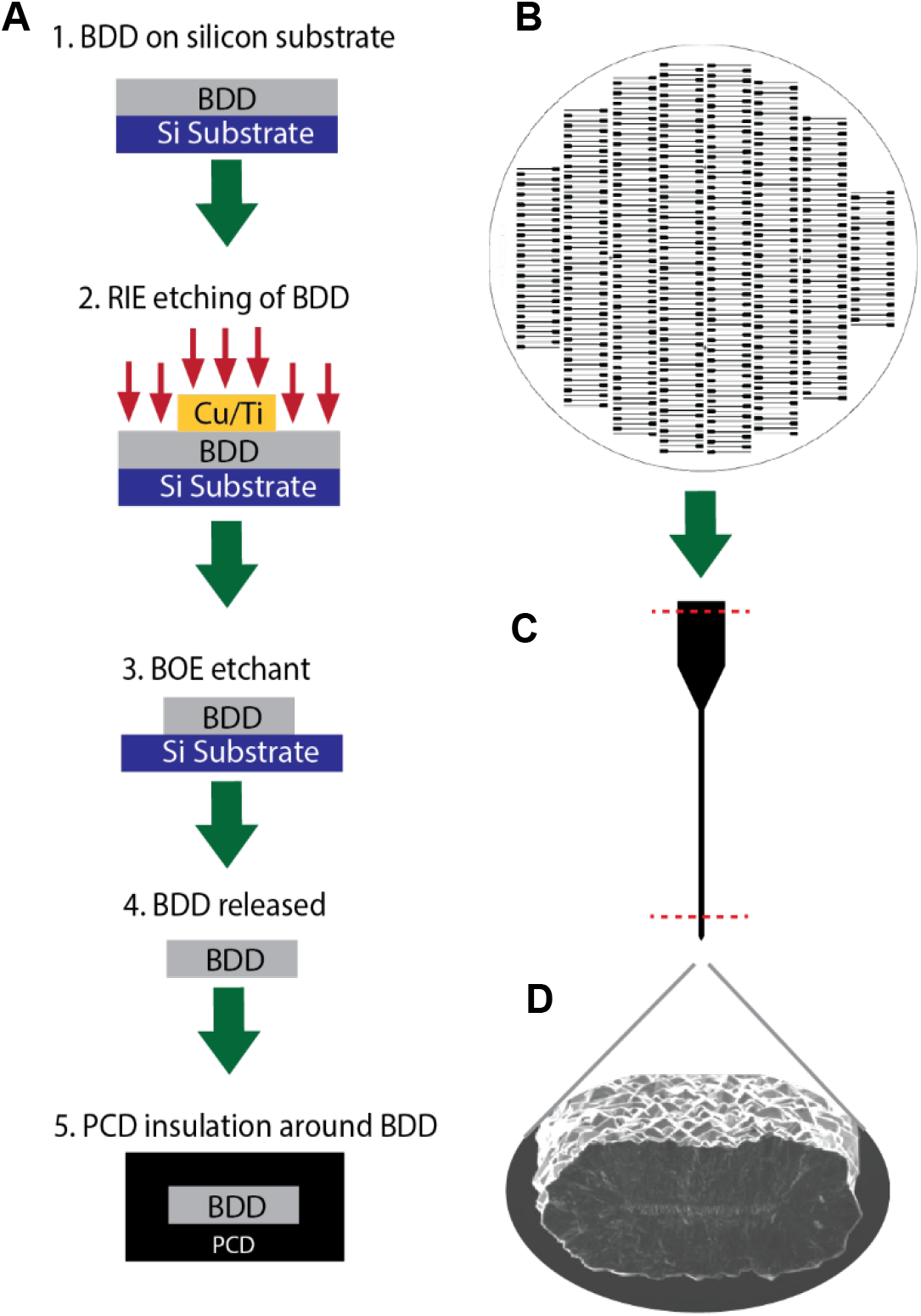
Fabrication of all diamond electrodes, where a conductive diamond core was insulated with an intrinsic insulating polycrystalline shell. **A**. Fabrication scheme of all diamond electrode where first boron doped diamond (BDD) is grown on a silicon wafer. Using lithography and a metal hard mask, the diamond is plasma dry etched and patterned to form the shape of the electrode. The silicon wafer was dissolved releasing the conductive core, and insulated by growing intrinsic polycrystalline diamond. **B**. Representative example of the scalability and flexibility of designing 100’s of electrodes simultaneously. **C**. Electrical connection is formed by cleaving the tip and connection pad. Dotted lines represent cleaving points to expose BDD conductive core. **D**. Scanning electron micrograph of the cleaved diamond tip, revealing the insulating polycrystalline diamond encasing the conductive boron doped diamond core.

Following fabrication, the ends of the electrodes were physically cleaved to expose the BDD core, revealing the pristine diamond (**Figure 1C**). Electrical connection through the diamond substrate was made by manually cleaving the opposite end, connection pads. The BDDME’s were secured in place, and electrically connected to a breakout board using an aqueous-based graphite conductive adhesive (Alfa Aesar, Ward Hill, MA). Following fabrication, a scanning electron micrograph of the BDDMEs revealed a BDD core to have a geometric surface area of either 105 µm^2^ or 175 µm^2^ (based on a 30 or 50 µm wide pattern, and 4 µm thick BDD growth thickness), and an insulating PCD shell of 11 µm to 20 µm thickness based on the distance tip was cleaved along the shank (**Figure 1D**). Thicknesses varied as the growth rate increases with distance from the filaments used during the diamond growth.

## Results and Discussion

Following fabrication, we utilized cyclic voltammetry to characterize the electrochemical response of the BDDME to ruthenium hexamine (RuHex) (**Figure 2A**), ferrocene carboxylic acid (FcCOOH) (**Figure 2B**), and dopamine (DA) (**Figure 2C**), in Tris aCSF pH 7.4 from 10 to 500 mV/s. We additionally measured DA in H_2_SO_4_ (**Figure SI-1A**). Utilizing the Randle-Ševčík equation we calculated the electroactive area for each electrode, and generated a Tafel plot to calculate the heterogeneous rate constant for these materials^36,37^(**Table 1**). Interestingly, the average calculated heterogeneous rate constant for FcCOOH with the BDDME is comparable to values previously reported by Jarošová et al. for BDD based microelectrode measurements^38^. It should be noted that Jarošová et al. electrodes were fabricated through growing BDD on Pt, and as such would have a lower resistance as compared to the BDD films, where diamond is used as the electrical contact. We observed a slight difference for RuHex, where the rate constant was slightly slower on our BDD. Notably, our CV current responses indicate a slightly resistive electrode or surface, with electrical impedance of 2.1 MΩ at 1 kHz when measured in Tris aCSF (**Figure SI-1B**). When comparing 1 mM 5-HT in Tris (**Figure SI-1C**), the scan rate at 100 mV/s revealed that oxidation was extremely slow, and it was not possible to calculate the electroactive area. As such k_o_ was not calculated. This could indicate an electrode-solution interface resistance, as when, using a 4-point probe after the conductive diamond growth, we measured the BDD and average resistivity to be 4.6 mΩ*cm ±1.4 mΩ*cm. All electrochemical measurements were performed on a cleaved diamond surfaces without any electrochemical pre-activation steps.

**Table 1:**
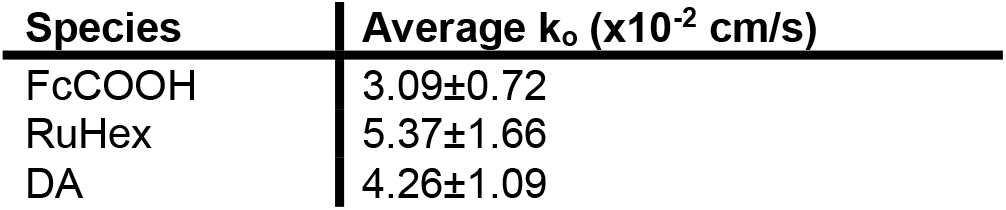
Heterogeneous rate constants for FeCOOH, RuHex and DA measured in Tris aCSF (pH 7.4) with different BDD µ-electrodes determined from Tafel Plots. (mean ± SEM, n=4 electrodes). All analyte concentrations were 1 mM.

| Species | Average $k_o$ ($\times 10^{-2}$ cm/s) |
| --- | --- |
| FcCOOH | 3.09±0.72 |
| RuHex | 5.37±1.66 |
| DA | 4.26±1.09 |

**Figure 2:**
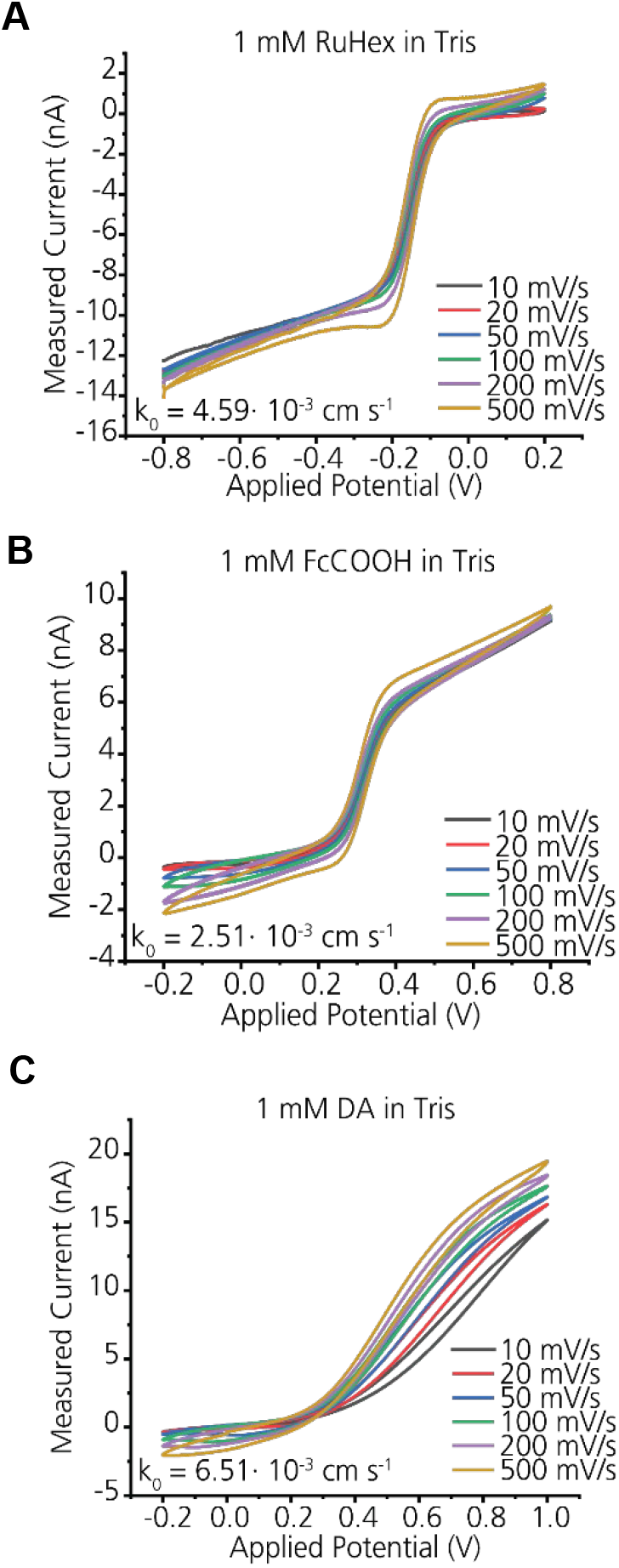
Slow scan cyclic voltammetry at the BDDME to determine electron transfer kinetics. **A**. Representative measurements of Ruhex on a BDDME, with the calculated rate constant for this electrode. **B**. Representative measurements of FcCOOH on a BDDME, with the calculated rate constant for this electrode. **C**. Representative measurements of DA on a BDDME, with the calculated rate constant for this electrode. All measurements were made in pH 7.4 Tris buffer. Applied potential was vs. reference electrode.

Following slow scan measurements, the BDDME was characterized in a custom flow cell injection system. The electrodes were pre-cycled utilizing the DA waveform applied at 10 Hz for 120 mins to study the background and electrode stability (**Figure SI-2**). The FSCV background current is generated by the rearrangement of ions around the electrode or charging of the electric double layer^39^. A high scan rate (400 V/s) captures the faradaic current produced by the redox reaction of electroactive species at the electrode^40,41^, but a large capacitive background current is also generated and needs to be subtracted out to identify the differential signal^40,42^. Thus, the stability of the background signal is pivotal for accurate analyte measurement. Previous literature has established the BDD electrode as an attractive electroanalytical detector for its low and stable background current over a wide working potential range^43–45^. Similarly, our all diamond BDDME demonstrated quick stabilization (within 15 minutes) and minimal drift over 120 min of continuous cycling of the electrode (**Figure SI-2A**). The background peak current on average was ∼22 nA at 0.6 V for a 30 µm wide electrode and ∼28 nA for a 50 µm wide electrode (**Figure SI-2C**). The shape maintained a low capacitive response. At the turn around potential, there was a slight peak increase. This is attributed to possible sp^2^ etching that occurs at the grain boundaries. We observed that during the initial cycle, the peak slightly decreased, however over the course of the 2-hr experiment, had minimal change. Interestingly on the back scan, there are non-capacitive changes, indicating slight electrochemical modifications to the diamond surface, which warrant further studies.

Previous publications have often focused on thin film, planar BDD electrodes, but limited studies have explored diamond as a microelectrode^29,46^. Park et al. (2005) reported on the background cyclic voltammetric response of thin film BDD on sharpened Pt wire in comparison with the carbon fiber microelectrode (CFME), and showed a low and stable background current independent of solution pH of the diamond microelectrode^47^. These BDD-Pt wire electrodes had a surface area of 1.8 × 10^4^ µm^2^ and were hydrogen terminated prior to electrochemical measurements to ensure pH insensitivity. However, hydrogen-termination of the electrode removes the surface-oxygen functionalities^48^ that are essential for adsorption controlled electrochemical measurements specifically for DA, using FSCV^49,50^. Our freestanding BDDME is unique in its micromachined fabrication process, smaller size (100 to 200 µm^2^), and bare, untreated surface favorable for FSCV measurements. At the same time, lack of hydrogen-termination of the our BDDME may leave some redox-active carbon-oxygen functionalities on the electrode surface, making the electrode sensitive to pH changes (**Figure SI-3**). It is also important to note that while our BDDME maintains a bulk sp^3^ carbon microstructure, some sp^2^ carbon may exist at the grain boundaries due to boron doping content. As shown in **Figure SI-2A**, the BDDME background was relatively stable over two hours with a slight shift that presents as a drift, possibly indicative of surface etching of the minor sp^2^ carbon present on the electrode surface. The BDDME showed excellent response stability and repeatability over this time with an average current response of 0.293 ± 0.007 nA for 5 µM DA injections (**Figure SI-2B**). The stability of the electrode over the course of two hours, and the stability of measuring ferrocene carboxylic acid (**Figure SI-2D**), demonstrates that the BDDME, without the metal support, maintains physical qualities that allow good chemical and electrical properties for an electrochemical sensor.

Fast stabilization of the all diamond BDDME can be especially advantageous for in vivo measurements where stabilizing an electrode immediately upon implantation can take long periods of time (up to 1 hour) before a single recording. When cycled in a conductive aqueous solution with an applied potential over 1.0 V, a carbon electrode surface can undergo chemical and physical changes that manifest as background drifts^42,51^. It has been reported that the BDD electrode lacks the majority of sp^2^ carbon structure and π bond network that favors such surface changes^29^, and thus, the background reaches equilibrium relatively quickly with minimal shifts. Our BDDME similarly contains low levels of sp^2^ carbon content, mainly concentrated at the grain boundaries, allowing the electrode to reach fast equilibrium for adsorption-controlled electrochemical measurements. (**Figure SI-4**) The Raman spectra showcases a strong boron doping peak at 491 cm^-1^, a diamond peak at 1210 cm^-1^, a disorder band at 1325 cm^-1^, and a small graphitic (G) band at 1540 cm^-1 52^.

To characterize the response of these diamond sensors and determine the diamonds sensitivity towards common neurotransmitters, we initially characterized the response of DA and 5-HT, AA, DOPAC, and H_2_O_2_ on the BDDME. (**Figure 3** and **Figure SI-5**). Interestingly, all tested analytes maintained a linear response with concentration and were well resolved on the BDDME with a sharp on and off current response ensuring high fidelity of temporal resolution and minimized electrode chemical fouling. A wide sensitivity difference between DA and 5-HT for the same concentration of each analyte (representative 10 µM response of DA and 5-HT shown in **Figure 3A&B** respectively) was observed. The electrode showed a linear current response to increasing concentrations of DA (**Figure 3C**) over the measured range of 0.2 to 10 µM with a limit of detection (LOD) of 0.26 µM found from the linear dynamic range of 0.2 to 10 µM. Similarly, a linear current response to increasing concentrations of 5-HT (**Figure 3D**) was observed over the same range of 0.2 to 10 µM, but with a lower limit of detection (LOD) of 0.16 µM calculated from the linear dynamic range of 0.2 to 2.0 µM (summary of statistics is reported in **Table 2**).

**Table 2:** Comparison for LOD and sensitivity for dopamine and serotonin using the BDDME (n=3) with an electroactive area of ∼100 to 200 µm^2^.

| Analyte | LOD ($\mu\text{M}$)* | Slope ($\text{nA}\mu\text{M}^{-1}$) | Measured Range ($\mu\text{M}$) | $r^2$ |
| --- | --- | --- | --- | --- |
| Dopamine (DA) | 0.26 | 0.046 | 0.2 – 10.0 | 0.999 |
| Serotonin (5-HT) | 0.16 | 0.192 | 0.2 – 10.0 | 0.996 |

\*Limit of detection (LOD) was calculated for the linear dynamic ranges of 0.2–10 $\mu\text{M}$ for DA and 0.2–2.0 $\mu\text{M}$ for 5-HT. See methods section for LOD calculation.

**Figure 3.**
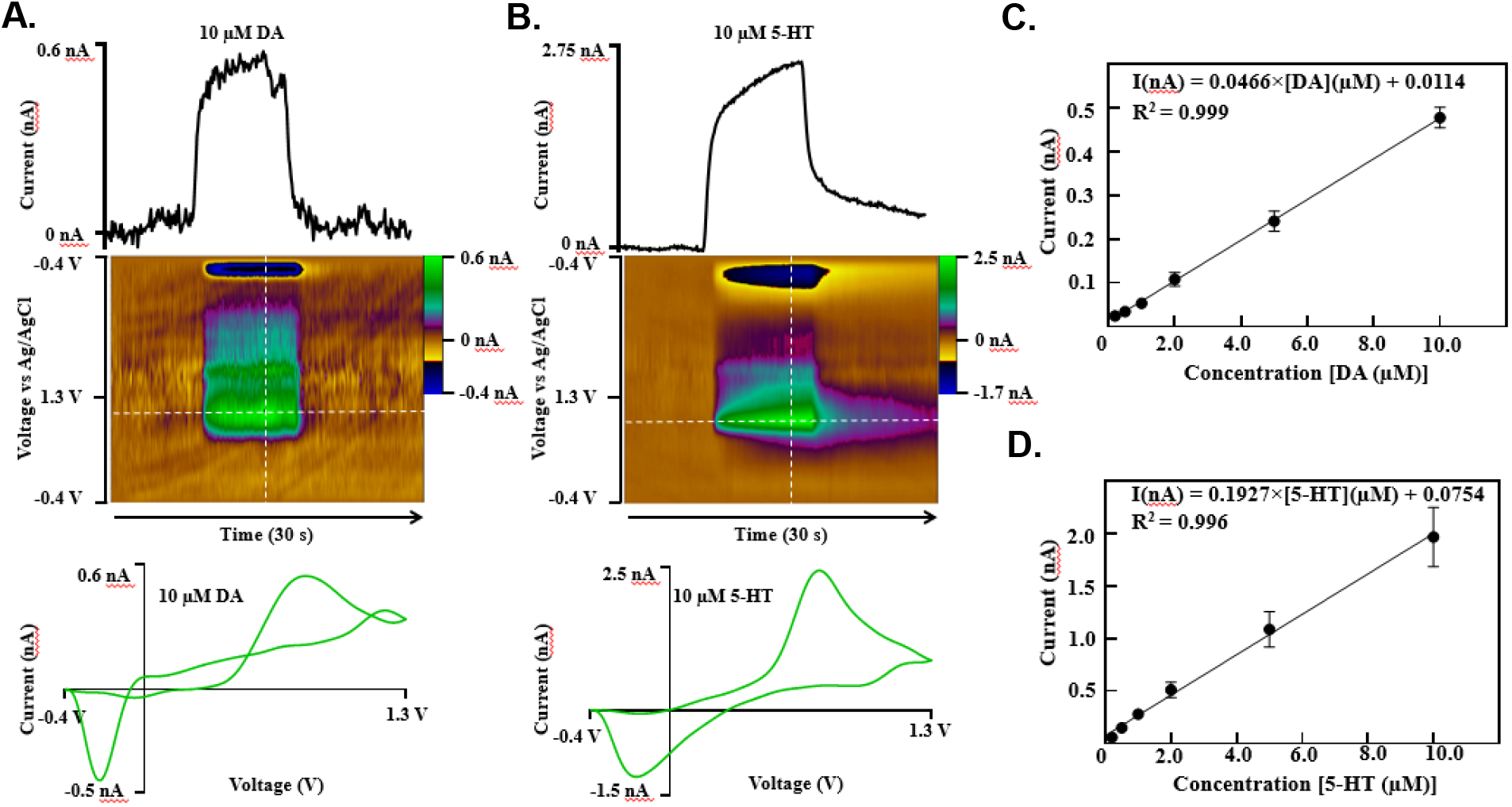
Comparison of the response for DA and 5-HT for the BDDME. The triangular waveform from -0.4 V to 1.3 V and back at 400 V/s was applied at 10 Hz frequency for all measurements. **A**. Representative FSCV current vs time trace, voltage vs time color plot, and current vs voltage cyclic voltammogram for 10 µM DA at the BDDME. **B**. Representative FSCV current vs time trace, voltage vs time color plot, and current vs voltage cyclic voltammogram for 10 µM 5-HT at the BDDME. **C**. Calibration curve of DA at the BDDME (*n*=3). **D**. Calibration curve of 5-HT at the BDDME (*n*=3).

Both neurotransmitters generate cyclic voltammograms (CVs) with oxidation and reduction peaks (**Figure 3 A&B**), but 5-HT has a larger and clearer response than DA. Notably, the oxidation and reduction peaks are prominently not symmetrical for DA, and a smaller third peak is evident near the turnaround potential of 1.3 V. The small peak at 1.3 V is theorized to be background subtraction error due to slight changes in the background. The representative CV for DA on the BDDME (Current Voltage plot, **Figure 3A**) is similar to a more traditional and symmetric “duck-shaped” CV with equivalent oxidation and reduction magnitudes that would be expected with slow scan voltammetry where there is less mass transport of the analyte to the electrode surface due to the growing diffusion layer^40^. The shape indicates that diffusion-controlled kinetics are at play, and the current is unable to drop to zero before the potential is reversed to induce reduction of the analyte^40^. Additionally, the CV has an oxidation peak ∼0.7 nA, with a reduction peak close to -0.5 nA. Ideally, if adsorption-controlled kinetics were solely governing the DA redox reaction, the FSCV voltammogram would generate a reduction peak size that is approximately half the area of the oxidation peak due to strong adsorption of DA and quick desorption of dopamine-*o*-quinone, oxidized form of DA, at the surface^40,50^. Here, the peak heights are nearly the same indicating that possibly adsorption properties of the BDDME, although containing some sp^2^ carbon content, do not allow complete and strong adsorption of DA with fast desorption of dopamine-*o*-quinone. On the other hand, the CV for 5-HT (Current Voltage Plot, **Figure 3B**) presents a cleaner response with an oxidation peak of 2.5 nA and reduction peak of -1.5 nA. Still, even in the 5-HT response, the current does not drop to zero before the potential is reversed and a less prominent yet visible second peak is present near the turn around. Based on the calibration curves and compared statistics (**Table 2**), the BDDME is favorable to detect lower limits of 5-HT than DA over the measured range of concentrations. Overall, this means that the BDDME is able to carry out adsorption-controlled redox reactions for the most part, but diffusion-controlled kinetics interfere possibly due to the sp^3^ carbon content at the surface.

We further characterized these electrodes against DOPAC, and AA (**Figure SI5-A&B**) using the standard triangular waveform, which both maintained a linear response with concentration. Interestingly, for DOPAC, the oxidation response was only resolved for the applied current range, and occurred predominantly at +1.1 V. No reduction peak was observed, indicating that a lower applied holding period is necessary to resolve the response. For AA, the oxidation peak was clearly resolved. All samples for the calibration curve were pH adjusted, to account for decreases in pH due to AA concentration. Comparing DA to DOPAC, a 10 µm DA maintains a similar response to 100 µm. DOPAC with the same current magnitude response. Similarly, for AA, 50 µm of AA correlates with 10 µM DA maintaining sensitivity favorability towards 10 µM DA. Interestingly for H_2_O_2_ there is a much better sensitivity on the BDDME, where 1 nA of response is equivalent to 2 µM (**Figure SI5-C**). It should be noted that the H_2_O_2_ measurements were made with an increased potential of 1.5 V for the turnaround to resolve the oxidation. Key metrics and responses for the analytes are reported in **SI-Table 1**.

Comparing the all diamond cleaved BDDME towards others which we have previously developed and reported on, it should be important to note that these BDDME electrodes with a cleaved tip show lowered sensitivity towards DA. Our previously reported all diamond electrodes were prepared using an ND:YAG 1064 nm IR laser to cleave and create the recording site^15^. No long-term studies were done in this report to understand the impact of laser dicing and electrode stability. More recently, our findings using a different laser slicing system with an 800 nm femto cutting laser show-cased the temporary increase in DA and 5-HT sensitivity, however these electrodes lacked long term electrochemical stability due to rapid sp^2^ rapid etching induced by the laser slicing^34^. Additionally, it was found in both studies that laser slicing damaged the PCD layer, increasing the effective surface area of the electrode by an unknown, and minimally controlled amount. All pieces of work point towards the need for diamond surface modification to increase the sensitivity for DA and 5-HT without causing long-term changes, which may lead towards chemical etching of the electrode itself.

While the BDDME may present a surface with weaker adsorption than the standard bulk sp^2^ carbon fiber electrodes, advantages to a less adsorptive surface include: (1) resilience towards chemical and physical surface changes in an analyte rich environment such as the brain, and (2) distinguishable neurotransmitter peaks due to adsorption favorability of one analyte over the other. Poor chronic stability and biofouling, i.e. adsorption of interferent proteins upon implantation in a biological environment, are long-standing challenges that disrupt neurochemical detection in the brain. We have previously reported that the BDDME experienced significantly lower biofouling-induced current reductions for 5-HT when using the selective “Jackson” waveform in comparison with the highly adsorptive CFME^12^. This indicates that the BDDME could potentially serve as a more resilient electrode in a chronic in vivo setting. Another prevalent FSCV in vivo challenge is accurate identification of neurotransmitter peaks – analytes that oxidize and reduce in the same potential window may overlap or appear together, complicating selective detection of the neurotransmitter of interest. Based on our observations from Figure 3, we exposed the BDDME to boluses of 10 µM 5-HT and 10 µM DA, individually followed by mixtures using the flow injection system to test peak distinguishability. We made mixtures with 1:1 (10 : 10 µM) DA to 5-HT, 1:2 (10 : 20 µM) DA to 5-HT, and 2:1 DA to 5-HT (20 : 10 µM) to study the FSCV voltammograms and peak shifts (**Figure 4**). As expected, a higher concentration of 5-HT in the mixture overwhelmed the response, and reduced the oxidation and reduction peaks’ separation. A higher concentration of DA in the mixture pulled the peaks further apart, indicating slower electron transfer when DA is present in the mixture.

**Figure 4.**
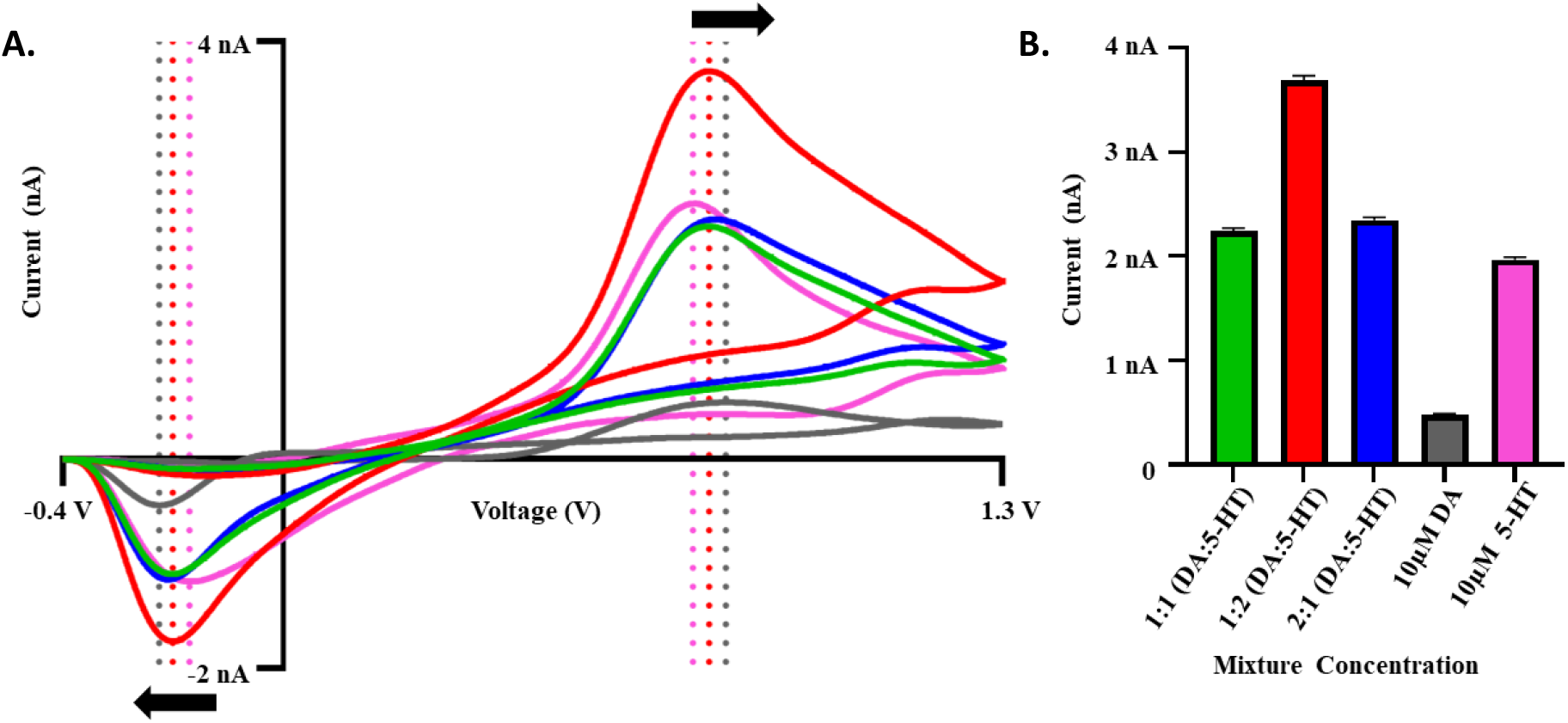
FSCV responses of mixed solutions containing DA and 5-HT on BDDMEs (*n*=3). **A**. Overlapping FSCV voltammograms of mixtures with varying concentrations of DA and 5-HT. Triangular waveform was employed, starting at -0.4 V to 1.3 V and back scanning at 400 V/s at 10 Hz frequency. Voltammograms of only 10 µm 5-HT and 10 µm DA plotted for reference. **B**. Corresponding current vs concentration plot comparing peak oxidative current for each solution. Mixtures (from left to right on x axis) contain 10 µm DA & 10 µm 5-HT (green), 10 µm DA & 20 µm 5-HT (red), 20 µm DA & 10 µm 5-HT (blue), only 10 µm DA (grey), and only 10 µm 5-HT (magenta).

The results in **Figure 4** show that the BDDME favors detection of 5-HT over DA even while using the standard triangular waveform employed for DA detection. There is a slight shift in oxidation response from 0.7 to 0.73 V when 5-HT is present as compared to the DA oxidation. There is an interesting additional change in the oxidation shape with DA present, where at 1.1 V, there is a broadening of the response. Additionally, there is a slight shift in the reduction potential as well from -0.35 to -0.3 V when 5-HT is present compared to DA. As there are no intrinsic differences in the oxidation measurements additional statistical methods such as PCA would need to be used to differentiate the two-chemical species. As both 5-HT and DA oxidize and reduce within the potential window of -0.4 V to 1.3 V, other manipulations to the waveform may need to be employed in vivo to selectively detect one neurotransmitter over the other. As previously reported, employing the Jackson waveform with a faster scan rate may increase the BDDME selectivity and eliminate the slower DA electron transfer, leaving a more prominent 5-HT signal. On the other hand, slowing the scan rate down may present a cleaner DA signal to match the sluggish electron transfer which is planned for future evaluations.

Various modifications to the electrode could further enhance electrochemical measurements at the BDDME, especially for in vivo applications. The BDDME reported here has an approximate geometrical electroactive surface area of ∼100 to 200 µm^2^, about ten times smaller than the standard carbon fiber microelectrodes (∼7 µm diameter, and 100 µm long for an area of 2238 µm^2^). Because the size of the detected current is proportional to the electroactive area on the surface of the electrode, increasing the electroactive area of the freestanding BDDME could contribute to greatly increasing peak current measurements and lowering the limits of detection. Additionally, as others for CFME have report on the effect of edge planes for DA adsorption^50^, diamond growth direction may impact chemical responses. Our measurements are made across the growth boundary, and previous research from our team has indicated a DA sensitivity difference between the growth face, and nucleation face of the BDD^26^. Alternatively, employing an array-style electrode could maximize detection sites and overall electroactive area. Laser cut BDDMEs, as reported in our recent study^34^, could provide a more sensitive, sp^2^ rich environment that may be beneficial for acute in vivo detection for neurotransmitter rich regions, however these are short term, as they etch due to the DA waveform switching potential of over 1.0 V. Pretreatment options with cation exchange polymers, such as Nafion, could also be advantageous for increased sensitivity of the electrode. These polymer films and membranes may increase adsorption limitations, as the diamond material would remain stable without etching from underneath the membrane, unlike the carbon fiber counterpart but remains to be studied.

## Conclusion

These findings showcase a scale-able all diamond probe and provide foundational information to guide future in vivo sensor development for electrochemical measurements using the freestanding BDDME. The results reported here are aimed at filling the knowledge gaps and background information to support our past and future work with the next generation of BDDME. The BDDME shows a favorability towards 5-HT detection with the oxidative peak responses with a 2-fold increase compared to DA for the same concentrations of analytes and a lower limit of detection of 0.16 µM for 5-HT than the 0.26 µM for DA. Similarly, the mixture comparisons elucidate that higher concentrations of 5-HT overwhelm the voltammogram, and DA demonstrates slower electron transfer in comparison to 5-HT at the BDDME surface. However, modifications to the waveform, and selective potential windows, may prove useful to differentiate between oxidative peak signals for the in vivo setting. Due to the surface characterization of the BDDME, a tradeoff in sensitivity may need to be accepted; however, this can be surmountable with increased electroactive area sizes, array-style electrode and surface treatments. These probes offer a biocompatible platform with stability in the measurement response for future consistent and chronic *in vivo* measurement of biomolecules.

## Acknowledgements

The authors would like to thank Robert Rechenberg for boron doped diamond growth, Davit Galstyan for intrinsic diamond growth, G. M. Hasan U. Banna for diamond lithography and structuring, and Michigan State University Center for Advance Microscopy for image analysis.

