## Supplemental Information for "All-Diamond Boron-Doped Microelectrodes for Neurochemical Sensing with Fast-Scan Cyclic Voltammetry"

### Supplemental Figures

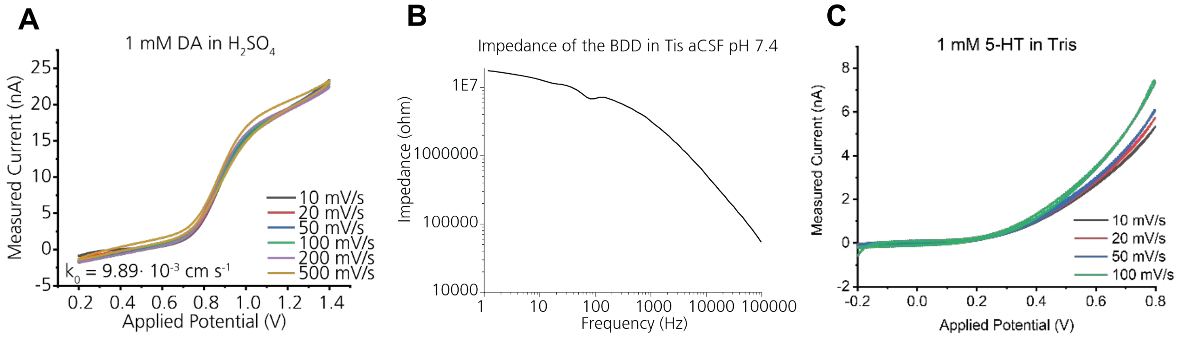


**Figure SI-1: A.** Voltammetric response of BDDME to DA in $H_2SO_4$. **B.** Representative impedance response of the BDDME in Tris aCSF. **C.** Representative 5-HT measurement in Tris aCSF solution.

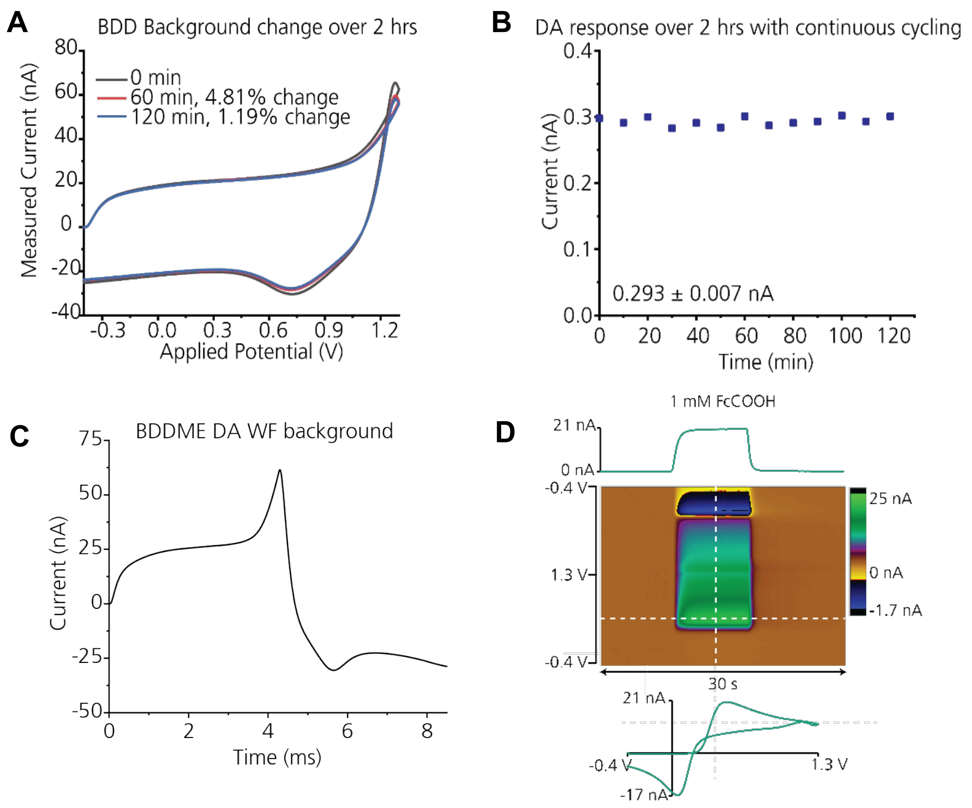


**Figure SI-2**: **A.** BDDME background response when applying the standard DA waveform at 10 Hz application frequency. **B.** Repeated injections of 5 μM DA delivered through the flow cell to the BDDME every 10 minutes showing a stable and reproducible response for the peak oxidative currents. **C**. Background of the BDDME with the triangular waveform commonly applied for dopamine detection starting at -0.4 V to 1.3 V and back at 400 V/s and at 10 Hz frequency. **D.** A representative response to measuring 1 mM FcCOOH in Tris pH 7.4, using a flow injection system on the BDDME.

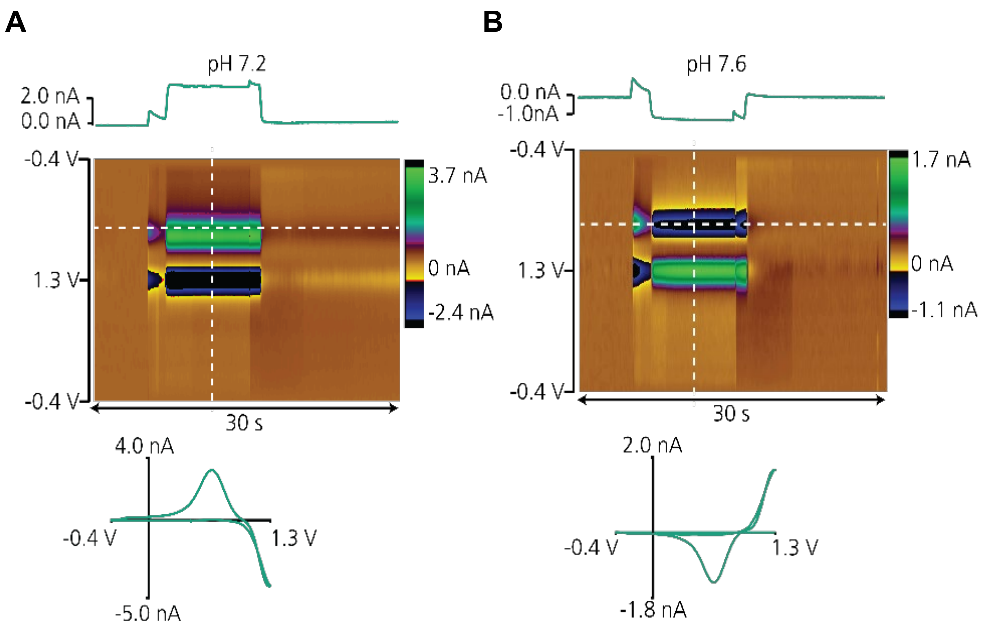


**Figure SI-3**. The BDDME was found to be highly responsive to pH changes, and the acidity of $HClO_4$ influenced the observed the current. The acidity of $HClO_4$ influenced the observed the current. Neurochemical stock solutions for in vitro experiments were in 1 mM $HClO_4$ as opposed to 0.1M $HClO_4$ that has been used previously reported in literature. High concentrations of $HClO_4$ overwhelmed the real DA current and presents on the backward scan.

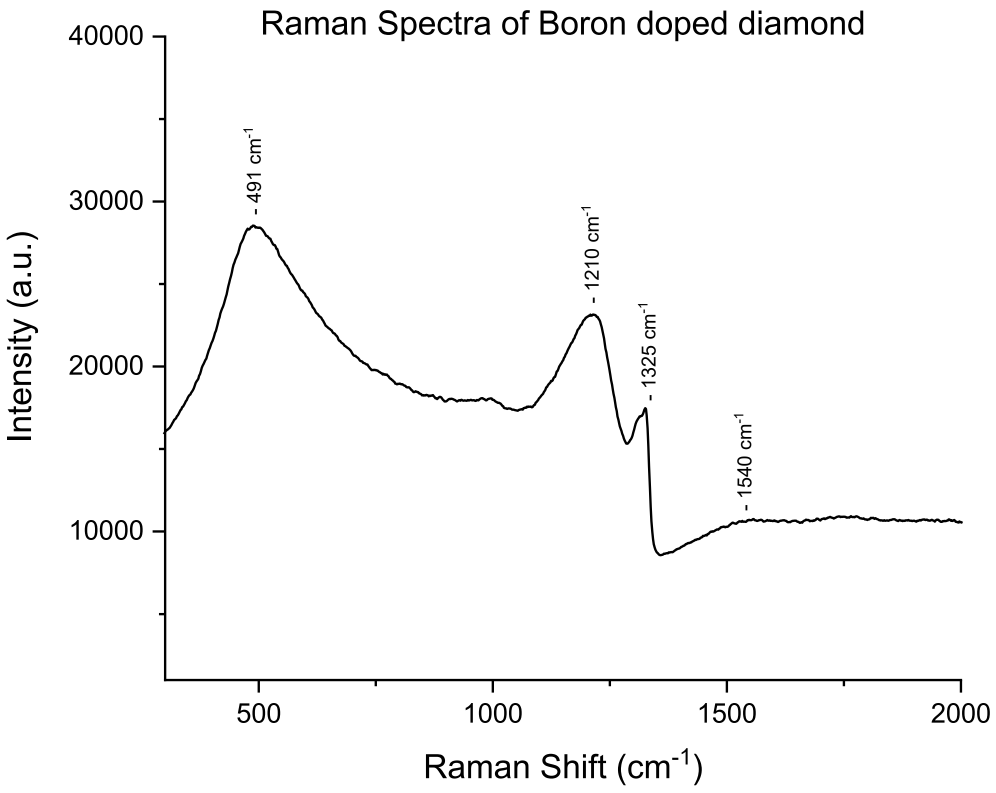


**Figure SI-4**. Representative Raman spectra for grown BDD. Representative Raman spectra of the conductive diamond used for the fabrication of the all diamond microelectrodes.

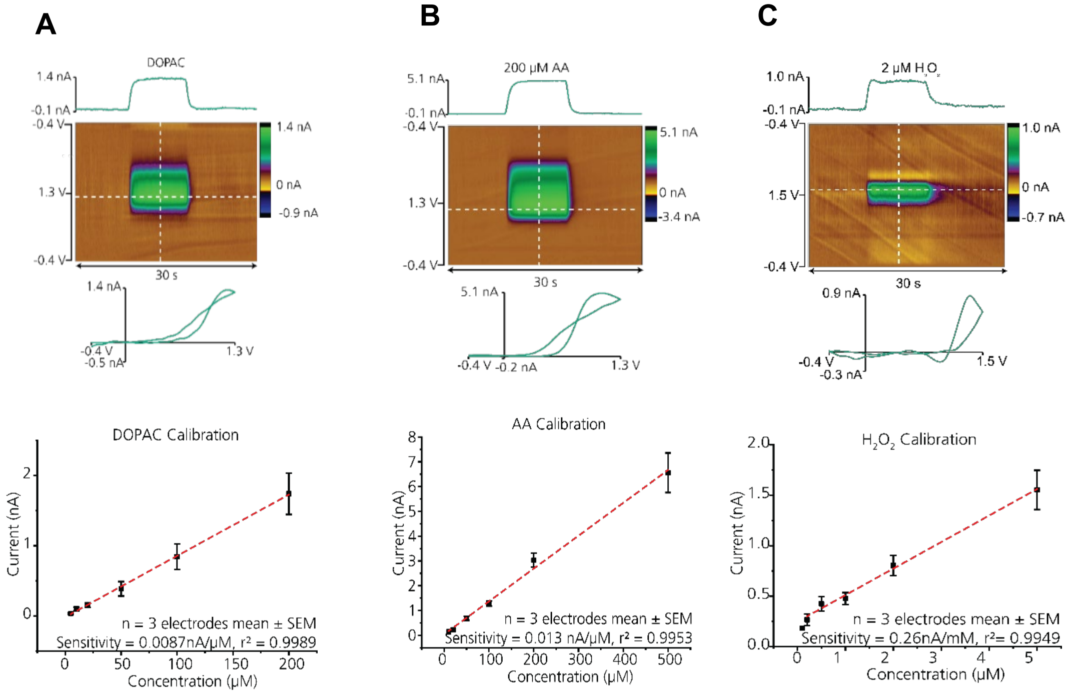


**Figure SI-5:** Commonly investigated NT responses at the BDDME. The triangular FSCV DA waveform from -0.4 V to 1.3 V to -0.4 V at a scan rate of 400 $Vs^{-1}$ with a frequency of 10 Hz was applied at the electrode for DOPAC and AA, and increased to 1.5 V for $H_2O_2$. A buffer solution of tris aCSF (pH 7.4) was pumped at a rate of 750 $\mu Lmin^{-1}$. Minimum of three buffer rinses were done prior to injecting analyte to flush the flow cell and prevent contamination.

**SI Table 1: Figures of merit of common compounds at the BDDME (electroactive surface area ~ 100 to 200 μm$^2$) (mean ± SEM, n=4 electrodes)**

| Compound | ΔEp (V) | Ox/Red ratio | Range of linearity (μM) | Slope (nA·μM$^{-1}$) | $r^2$ | Limit of detection (μM) |
|---|---|---|---|---|---|---|
| Ascorbic Acid | 0.67±0.08 | - | 5-200 | 0.00872 | 0.9989 | 3.26 |
| DOPAC (extended WF, -0.7-1.3 V) | 1.53±0.16 | 1.53±1.6 | 10-500 | 0.0133 | 0.9953 | 2.60 |
| pH | - | - | 7.2-7.6 (pH) | -5.52 (nA/pH) | 0.9856 | - |
| $H_2O_2$ (extended WF -0.4-1.5 V) | 1.40±0.01 | - | 0.1-5 | 0.263 | 0.9911 | 0.200 |
